# Pathogen-Host Analysis Tool (PHAT): an Integrative Platform to Analyze Pathogen-Host Relationships in Next-Generation Sequencing Data

**DOI:** 10.1101/178327

**Authors:** Christopher M. Gibb, Robert Jackson, Sabah Mohammed, Jinan Fiaidhi, Ingeborg Zehbe

## Abstract

**Summary:** The Pathogen-Host Analysis Tool (PHAT) is an application for processing and analyzing next-generation sequencing (NGS) data as it relates to relationships between pathogen and host organisms. Unlike custom scripts and tedious pipeline programming, PHAT provides an integrative platform encompassing raw and aligned sequence and reference file input, quality control (QC) reporting, alignment and variant calling, linear and circular alignment viewing, and graphical and tabular output. This novel tool aims to be user-friendly for life scientists studying diverse pathogen-host relationships.

**Availability and Implementation:** The project is publicly available on GitHub (https://github.com/chgibb/PHAT) and includes convenient installers, as well as portable and source versions, for both Windows and Linux (Debian and RedHat). Up-to-date documentation for PHAT, including user guides and development notes, can be found at https://chgibb.github.io/PHATDocs/. We encourage users and developers to provide feedback (error reporting, suggestions, and comments) using GitHub Issues.

**Contact:** Lead software developer

## Introduction

While the ease of producing and accessing next-generation sequencing (NGS) data has grown significantly, bottlenecks still exist in its processing and analysis. Alignment algorithms, particularly for short-read alignment and the tools that implement them, have matured to the point that they no longer present the major hurdle in the data analysis process (Li and Homer, 2010). The availability of fast and user-friendly tools has become the limiting factor (Milne *et al.*, 2010). There are excellent tools which perform one or several discrete functions in the same domain, *e.g.*, Bowtie2 (Langmead and Salzberg, 2012) and SAMtools (Li *et al.*, 2009), however all-in-one type platforms can offer a breadth of features that help address barrier-to-entry (*i.e.*, the ease in which users can setup and perform analyses). While integrative multi-tool platforms such as Comparative Genomics (CoGe) (Lyons and Freeling, 2008), VirBase (Li *et al.*, 2014), Pathogen-Host Interaction Data Integration and Analysis System (PHIDIAS) (Xiang *et al.*, 2007), Galaxy (Afgan *et al.*, 2016), and Unipro UGENE (Okonechnikov *et al.*, 2012) exist, all are server or cloud-based, or maintained by relatively small communities. The rather small scale of the infrastructure behind these projects and their cloud-based nature, introduce roadblocks in the transfer of data to and from their servers (Li and Homer, 2010). One solution to a problem such as this is to establish an on-site computational cluster, but that can add further barrier-to-entry for data analysis due to technical and infrastructure requirements.

We sought to develop the Pathogen-Host Analysis Tool (PHAT) to alleviate these issues by presenting an easy-to-setup and easy-to-use platform for life scientists conducting pathogen-host NGS analysis on common desktop computing hardware (*e.g.*, Windows).

## Features

Pathogen-host NGS analysis typically begins with high-throughput sequencing output files, containing experimentally relevant nucleic acid read information. PHAT provides a platform for analyzing these output data, with a focus on pathogen-host relationships (**Figure 1**). DNA read files are input into PHAT as FASTQ files (Cock *et al.*, 2010), comprised of sequence reads with per base nucleotide identities and quality scores, or pre-aligned SAM/BAM files (Li *et al.*, 2009) generated via cloud-based tools such as Galaxy (Afgan *et al.*, 2016). Quality control analysis can be performed on individual files, with graphical and numeric quality control reports generated. Reference genomes input as FASTA files must be indexed before they can be visualized or used for analysis. Once a pair of forward and reverse reads, paired FASTQ files, and a reference genome have been input, alignment of the paired reads against the reference can occur. PHAT also supports unpaired alignment and visualization of pre-aligned sequences.

**Figure 1.**
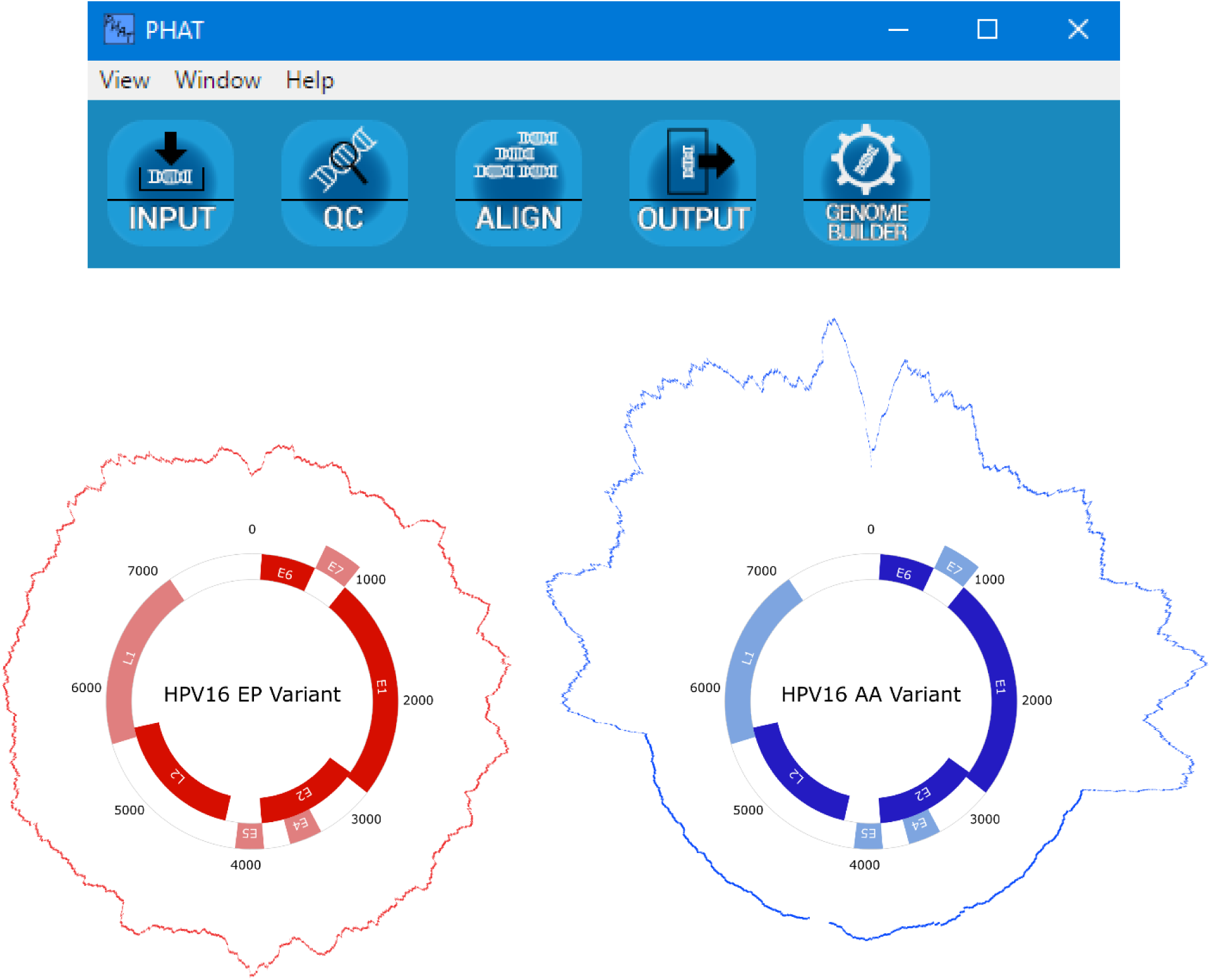
Pathogen-Host Analysis Tool (PHAT) tools and visualization. PHAT is equipped with tools (top image, showing the main toolbar) allowing the input of pathogen-containing next-generation sequencing data and reference sequences, quality control testing with FastQC, alignment with Bowtie2, SNP detection with VarScan 2, visualization with pileup.js and PHAT’s genome builder, as well as tabular output. Example human papillomavirus type 16 (HPV16) genome maps and coverage plots were generated with PHAT to contrast coverage between viral variants: the European Prototype (EP) variant is episomal, whereas the Asian-American (AA) variant’s coverage is disrupted by integration into host DNA. Data from Jackson *et al.*, 2016.

The core functions of the PHAT platform as well as FASTQ quality control, sequence alignment, and its automated analyses are performed through well-known, established implementations. Namely, quality control scoring for high-throughput sequencing data is performed by FastQC (https://www.bioinformatics.babraham.ac.uk/projects/fastqc/), sequence alignment by Bowtie2 (Langmead and Salzberg, 2012), alignment visualization by pileup.js (Vanderkam *et al.*, 2016), circular genome visualization based on our enhancements to AngularPlasmid (http://angularplasmid.vixis.com/) which we make available as a new project called ngPlasmid (https://github.com/chgibb/ngPlasmid), and automated variant calling by VarScan 2 (Koboldt *et al.*, 2012). The graphical user interface of the application, based on GitHub’s Electron project (http://electron.atom.io/docs/) operates in a client-server based architecture. Each window of the application acts as a client, communicating with a background server process. The server process manages the saving of and propagation of workspace data. This server process also manages the spawning of other processes used in the application such as sequence alignment and quality control report generation. This mechanism allows other processes to act as threads, allowing the flow of data to and from the application window that invoked it and the spawned process itself. On systems with limited power, the server process can limit the number of concurrently running processes as well as the amount of data propagated between application windows to reduce memory and CPU usage. Application windows which use other processes for operations or generation of data (*e.g.*, Bowtie2) utilize a pipeline, requesting the spawning of new processes as others close and passing data from process-to-process to arrive at the result (*e.g.*, the conversion of data from BAM format, to a sorted, indexed BAM). The server process, as well as the application windows themselves are implemented in TypeScript. These application windows can be conveniently undocked from the main toolbar as per user requirements.

## Conclusion

With the development of PHAT, we aim to bring simple-to-use, cross-platform NGS analysis to off-the-shelf hardware for life scientists studying pathogen-host relationships. In our own lab, we study human papillomavirus type 16 (HPV16) variants and their tumourigenicity in human skin using NGS (Jackson *et al.*, 2016), but PHAT can be applied to a wide-variety of pathogen-host relationships (*e.g.*, genotyping of microbes such as viruses, bacteria, and protozoans from host NGS samples). We plan to actively develop, update, and support PHAT, with auto-updating features already included, in anticipation of building an active user and developer community.

## Authors’ Contributions

The need for PHAT was initially conceived by Robert Jackson (RJ) and Ingeborg Zehbe (IZ). User interface (UI) and functional design was carried out by Christopher M. Gibb (CMG) and RJ collaboratively. Programming and technical planning was carried out by CMG. Manuscript and documentation writing and editing was carried out by CMG, RJ, Sabah Mohammed (SM), Jinan Fiaidhi (JF) and IZ. Intellectual property considerations made by SM and JF. Thanks to Zehbe lab members for user testing as well as students Mitchell Pynn, Jeremy Braun, Shane Liu, Zachary Moorman, and Nicholas Catanzaro for volunteering their time to technical improvements. We are thankful to the third-party developers of open-source tools that are incorporated in PHAT.

